# Rates of gene conversions between *Escherichia coli* ribosomal operons

**DOI:** 10.1101/2020.08.03.235192

**Authors:** Isaac Gifford, Aurko Dasgupta, Jeffrey E. Barrick

## Abstract

Due to their universal presence and high sequence conservation, rRNA sequences are used widely in phylogenetics for inferring evolutionary relationships between microbes and in metagenomics for analyzing the composition of microbial communities. Most microbial genomes encode multiple copies of ribosomal RNA (rRNA) genes to supply cells with sufficient capacity for protein synthesis. These copies typically undergo concerted evolution that keeps their sequences identical, or nearly so, due to gene conversion, a type of intragenomic recombination that changes one copy of a homologous sequence to exactly match another. Widely varying rates of rRNA gene conversion have previously been estimated by comparative genomics methods and using genetic reporter assays. To more directly measure rates of rRNA intragenomic recombination, we sequenced the seven *Escherichia coli* rRNA operons in 15 lineages of cells that were evolved for ~13,750 generations with frequent single-cell bottlenecks that reduce the effects of selection. We identified 34 gene conversion events and estimate an overall rate of intragenomic recombination events between rRNA copies of 3.2 × 10^−4^ per generation or 5.3 × 10^−5^ per potential donor sequence. This rate varied only slightly from random expectations between different portions of the rRNA genes and between rRNA operons located at different locations in the genome. This accurate estimate of the rate of rRNA gene conversions fills a gap in our quantitative understanding of how ribosomal sequences and other multicopy elements diversify and homogenize during microbial genome evolution.

## INTRODUCTION

Ribosomes perform some of the most highly conserved chemistry in cells—the translation of proteins from messenger RNAs—and ribosomal RNA genes are the most conserved genes across all domains of life (Isenbarger *et al.* 2008). Because of their low rate of evolution and ubiquity across taxonomic divisions, the nucleotide sequence of the small subunit RNA, known in bacteria and archaea as the 16S and in eukaryotes as the 18S, is often used to measure divergence between distantly related species (Woese 1987) and is therefore one basis for reconstructing the “Tree of Life” (Fournier and Gogarten 2010). Small ribosomal subunit amplicon sequencing is also commonly used to estimate the representation of different species in environmental and host-associated microbial communities (Okuda *et al.* 2012; Langille *et al.* 2013).

The need for new protein synthesis is often the dominant factor limiting cellular replication (Scott *et al.* 2010; Ceroni *et al.* 2015), and ribosomes can account for up to 20% of the dry mass of rapidly dividing bacterial cells (Weider *et al.* 2005). To meet this high demand, most bacterial genomes encode multiple ribosomal rRNA operons (Roller *et al.* 2016) and rRNAs are transcribed from highly active promoters (Paul 1994). Bacterial species capable of more rapid growth generally have more rRNA operon copies in their genomes (Klappenbach *et al.* 2000; Roller *et al.* 2016), and deleting some of these operons generally reduces the maximum growth rate that a bacterial strain can sustain (Asai *et al.* 1999, Stevenson and Schmidt 2004).

Ribosomal RNA genes within a single organism tend to have extremely similar, if not identical, sequences. This is thought to be due to a high rate of gene conversion, a type of non-reciprocal intragenomic homologous recombination between highly similar sequences that converts one into a precise copy of the other (Liao 2000). In bacteria, gene conversions occur through RecA-dependent recombination (Prammananan *et al.* 1999) and can occur with or without crossing over (Santoyo and Romero 2004). Gene conversions can be either allelic, within multiple copies of the same gene, or ectopic, between sufficiently similar genes (Galtier 2001). Allelic gene conversions can lead to concerted evolution of multicopy genes when it occurs at a sufficient rate that new mutations appearing in one copy are likely to be either reverted or propagated to all of the other homologous loci (Liao 1999, Liao 2000, Gogarten *et al.* 2002). Since they homogenize multicopy genes, allelic gene conversions have been proposed to be important for maintaining the ability of the cell to interchangeably use all copies of these genes for assembly of complex, multiple-subunit structures, like ribosomes (Liao 1999).

Accurate estimates of gene conversion rates are necessary to understand how they interact with other mutational processes during genome evolution. However, current estimates of the frequencies of gene conversions between *Escherichia coli* rRNA operons range over many orders of magnitude (Harvey and Hill 1990; Hashimoto *et al.* 2003). These studies, like many others looking at gene conversion, rely on reporter assays in which a single type of gene conversion is observed because it leads to a new phenotype that has a strong effect on cellular fitness (restored growth rate or antibiotic resistance, respectively). In such cases, viewing these frequencies through the prism of natural selection skews the inferred rates of gene conversions. By contrast, microbial mutation accumulation (MA) experiments propagate populations for many generations through single-cell bottlenecks to eliminate selection against all but the most deleterious mutations (Halligan and Keightley 2009). Mutation rates can be accurately determined from MA experiments simply by counting how many genomic changes of different types appear over time in replicate lineages (Foster et al. 2015).

We previously analyzed mutation rates in a mutation accumulation experiment in which independent lineages of *Escherichia coli* B were propagated for 550 days (Kibota and Lynch 1996; Tenaillon *et al.* 2016). Due to limitations of short-read resequencing data, we could not measure the rates of gene conversions in rRNA operons with that data. Here, we sequenced heterologous sites in all seven copies of the 16S and 23S genes in 15 endpoint strains from the MA experiment. We use the observed changes to estimate an overall rate for gene conversions between highly homologous sequences and examine whether this rate varies substantially between different copies of the rRNA genes and sites within them.

## MATERIALS AND METHODS

### rRNA operon sequencing

The MA experiment has been described previously (Kibota and Lynch 1996; Tenaillon *et al.* 2016). Briefly, MA lineages were started from a 2,000-generation clone isolated from a long-term evolution experiment (REL1206) that differs by a few mutations from *Escherichia coli* B REL606 (Jeong *et al.* 2009). The MA experiment consisted of daily transfers in which a randomly selected colony was streaked out on Davis minimal medium agar supplemented with 200 μg/ml glucose. An estimated 13,750 generations elapsed over the course of the 550-day experiment (Tenaillon *et al.* 2016). For amplification, we divided each of the seven *E. coli* rRNA operons into two PCR amplicons, one covering the *rrs* gene (16S) and the other including the *rrl* gene (23S) and the first downstream *rrf* gene copy (5S). PCR products for REL1206 and endpoint clones from 15 MA lineages were sequenced with the primers used for amplification and additional internal primers. Sequencing covering all heterologous sites was carried out at the UT Austin DNA Sequencing Facility on an ABI 3730xl DNA Analyzer (Thermo Fisher).

### Gene conversion inference

Trace files from PCR amplicon sequencing were aligned to the LTEE ancestor (REL 606) using Geneious 6. We found that REL1206 differed from REL606 by a single gene conversion: the 23S-5S spacer in the *rrnD* operon was converted to the *rrnAEH* type. In all instances in which a particular rRNA copy in the evolved stain differed from the corresponding copy in REL1206, the changes exactly matched at least one of the other rRNA copies. We inferred the properties of the most parsimonious set of contiguous gene conversions that could lead to these changes in sequence as follows. First, we determined which rRNA copies could have been donors for all bases changed by a given conversion. Next, we set the minimum conversion size as the length from the first changed base to the last changed base. Finally, we set the maximum conversion size as the largest 16S or 23S region from any of the donors that would only result in the observed changes. That is, we extended both ends through matching donor sequences until encountering a difference in all possible donor copies or the end of the rRNA subunit gene.

### Gene conversion rate analysis

The number of mutations of a given type that accumulate during an MA experiment is expected to follow a Poisson distribution if reversions occur at a negligible rate and opportunities for observing new mutations do not appreciably decline over time. Based on the small overall numbers of base changes in rRNA operons that we observed in each MA lineage, these assumptions should largely hold in our dataset. Therefore, we estimated rates and confidence limits on those rates from gene conversion counts using Poisson distributions with a total time basis of 206,250 generations (15 independent lineages × 13,750 cell doublings).

An additional consideration is that many gene conversion events between rRNA operons will be undetectable because they occur between two sequences that are already identical. In order to correct our rate estimates for these unobservable gene conversions we performed bootstrap resampling using a custom Python script (**File S1**). This procedure utilized an alignment of the 16S, 23S, and 5S rRNA sequences from the REL606 genome (Jeong *et al.* 2009) aligned with MUSCLE v3.8.31 (Edgar 2004). We generated sets of random conversions by selecting random sizes, locations that placed these regions entirely within the rRNA subunit alignment, and donor and recipient operons. Conversion sizes were determined by choosing a random conversion event from the observed data with replacement and then choosing a random size between the minimum and maximum inferred sizes for the selected conversion. A minimum size of 50 bp was enforced for small conversions because this is the required size for efficient RecA-mediated homologous recombination in *E. coli* (Singer *et al.* 1984; Watt *et al.* 1985). Using larger minimum conversion sizes of 100-300 bp did not appreciably alter our results. Finally, donor and recipient operon sequences in the selected region were compared to determine if a change of sequence would be observed from that resampled conversion.

To estimate gene conversion rates, we performed sets of 10,000 replicates in which we drew a total number of gene conversion events from the Poisson distribution with a candidate rate and used the resampling procedure to determine how many of these resulted in observable changes. The maximum likelihood estimate of the gene conversion rate per genome was determined as the rate with the highest probability of resulting in exactly the actual number of observed gene conversions. The 95% confidence interval on this estimate was determined by finding a lower rate with a 2.5% tail probability of resulting in the number of observed gene conversions or more and a greater rate with a 2.5% tail probability of resulting in the observed number or fewer. To understand variation in the rates of gene conversions between different rRNA operon copies and within different portions of their sequences, we performed 10,000 replicates of the resampling procedure in which we continued to draw new conversions until there were exactly as many conversions resulting in sequences changes as were empirically observed. This allowed us to estimate 95% confidence intervals on the number of each type of change by taking the corresponding 2.5% and 97.5% quantiles from the 10,000 sets of resampled gene conversions. The *p*-values for observed conversions for each operon or site relative to the randomized datasets were calculated as twice the proportion of replicates with values as or more extreme than the observed conversion numbers and adjusted for multiple testing with a Bonferroni correction.

We considered the four gene conversions involving the 23S-5S rRNA spacer separately because of the lower sequence homology in this region compared to positions within the 16S and 23S rRNA copies. There are two sequence variants of the 23S-5S rRNA spacer in the MA ancestor strain REL1206. Because they are present in 3 and 4 rRNA copies apiece, 24 of the 42 possible gene conversions between pairs of rRNA operons result in observable sequence changes. Therefore, we estimated a maximum likelihood conversion rate and corresponding exact 95% confidence intervals from the four observed events and multiplied each by 42/24.

### Changes in rRNA sequence identity

Concatenated sequences of the 16S, 23S, and 23S-5S rRNA spacer were used to compare the homology of sequences in each MA endpoint clone with the ancestor strain. Average percent sequence identity values were calculated as described by May (2004) using alignment length as the denominator for each pairwise combination of operons within each genome.

### Data Availability

Strains are available upon request. Supplemental files are available at FigShare. File S1 contains a Python script used to estimate the total gene conversion rate from observed gene conversions. File S2 contains alignments of the *rrs* and *rrl* genes and the *rrl*-*rrf* intergenic space sequences.

Table S1 contains gene conversions observed in the MA experiment.

## RESULTS AND DISCUSSION

The *E. coli* chromosome encodes seven ribosomal operons, with a majority located near the origin of replication (**Fig. 1a**). We sequenced amplicons corresponding to the 16S (*rrs*) and 23S+5S (*rrl*+*rrf*) portions of each rRNA operon (**Fig. 1b**) in endpoint clones from 15 independently evolved *E. coli* lineages from a 550-day mutation accumulation (MA) experiment. We identified a total of 56 base substitutions and insertions or deletions of a few bases in the 16S and 23S genes of the evolved strains. All of these changes could be explained by gene conversions (as shown for one example in **Fig. 2**). We did not find any new alleles in the rRNA genes that would require *de novo* point mutations to explain. We also observed four conversions that switched the 23S-5S intergenic region between a 186-bp sequence that is initially present in four of the rRNA operons and a shorter 92-bp variant present in the other three.

**Figure 1.**
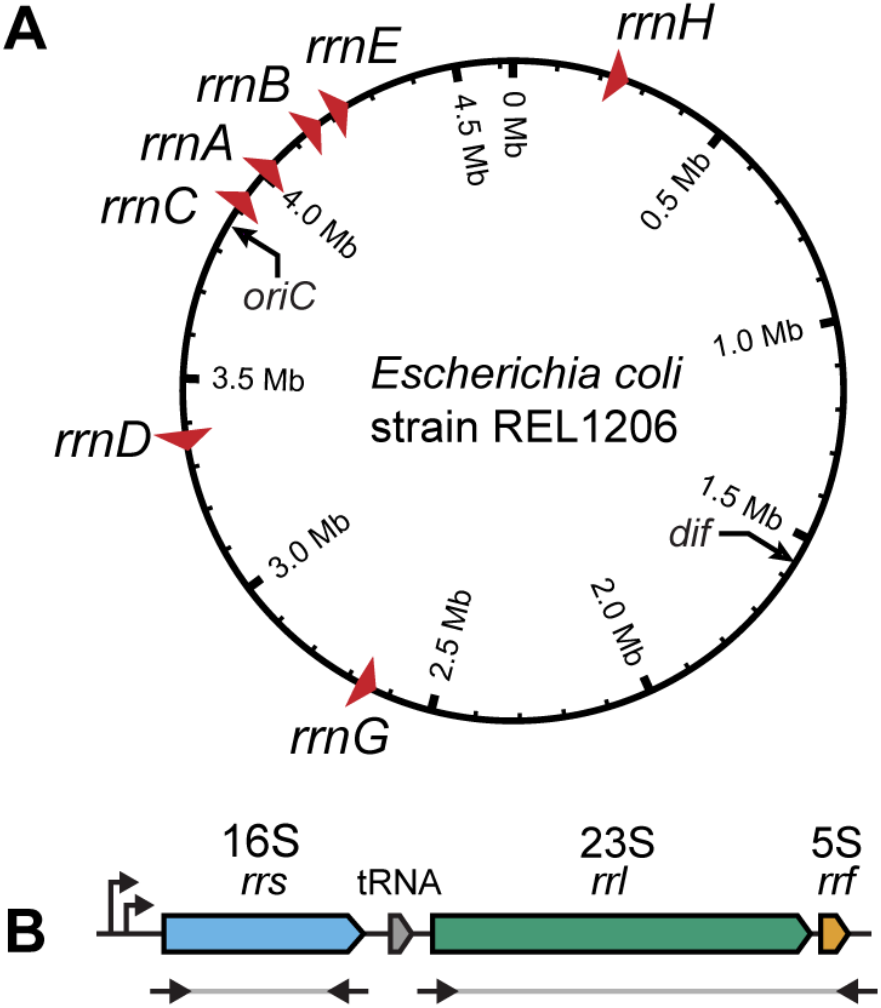
Ribosomal RNA operons in *E. coli*. (A) *E. coli* B strain REL606 chromosome showing the locations and orientations of the seven rRNA operons. (B) Organization of a typical rRNA operon showing the two PCR amplicons that were sequenced at heterologous sites in evolved genomes isolated at the end of a 550-day mutation accumulation experiment.

**Figure 2.**
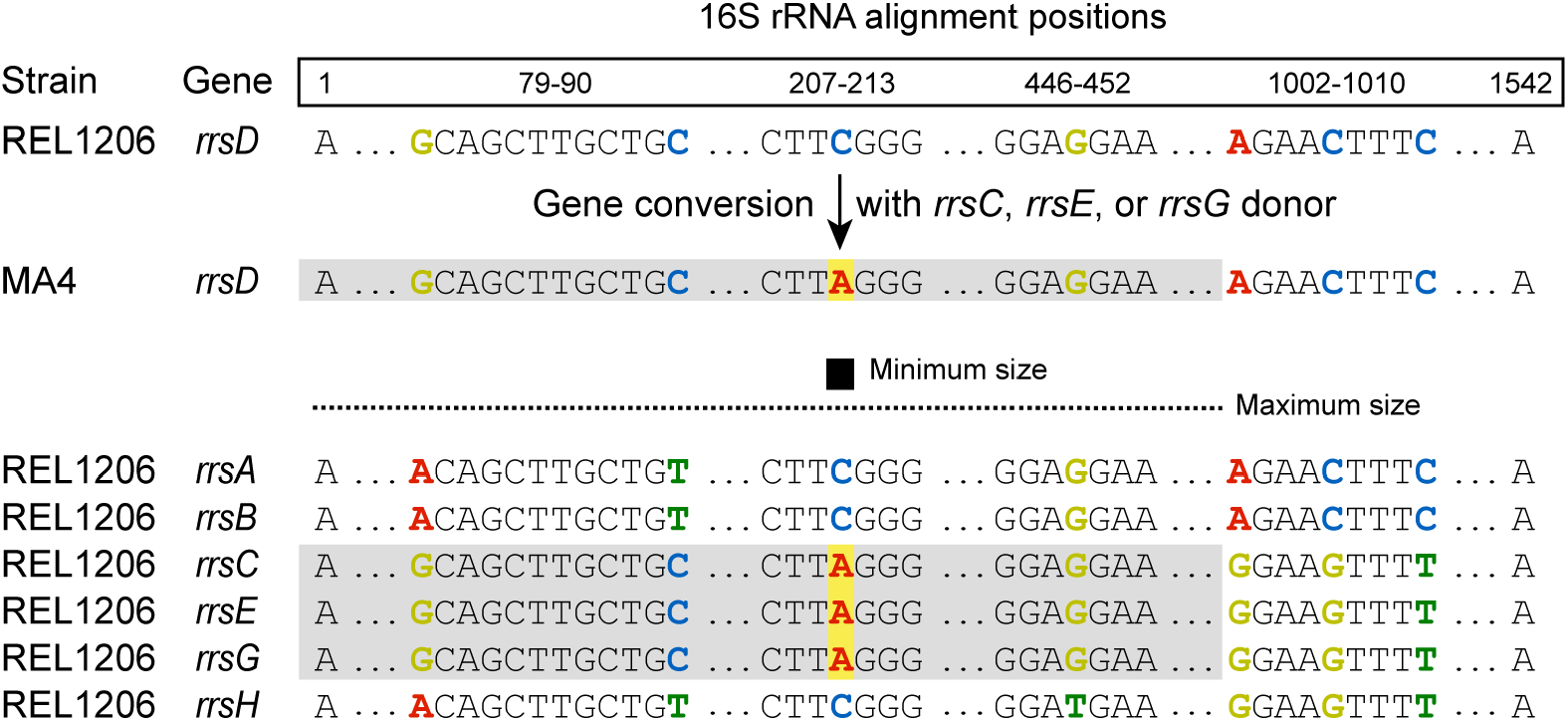
Example of a 16S rRNA gene conversion. In the sequenced endpoint clone from line 4 of the mutation accumulation experiment a change of a C to an A was observed at position 210 of the 16S subunit alignment in the *rrnD* operon. Either the *rrnC*, the *rrnE*, or the *rrnG* operon 16S sequence could have acted as a donor to cause this change through a gene conversion. Because the sequence of *rrnD* already matched all three of these donors at other alignment positions from 1-1001, the actual gene conversion could have been as large as 1001 base pairs.

The most parsimonious model that can reproduce the evolved rRNA sequences requires 38 gene conversion events: 7 in the 16S subunit, 27 in the 23S subunit, and 4 in the 23S-5S spacer (**Fig. 3, Table S1**). These conversions change anywhere from a single base to all bases that differ between two rRNA copies within a contiguous 2137-bp region. We calculated conversion rates of 5.8 × 10^−5^ (95% confidence interval: 2.3 × 10^−5^ to 1.2 × 10^−4^) and 2.6 × 10^−4^ (95% confidence interval: 1.7 × 10^−4^ to 3.8 × 10^−4^) per genome for the 16S and 23S gene conversions respectively. Though there is evidence that these rates are different from one another (likelihood ratio test, p =0.027), they were similar enough that we combined these observations to calculate an overall rate for these conversions that involve changing a small number of bases within mostly homologous sequences of 3.2 × 10^−4^ per genome (95% confidence interval: 2.2 × 10^−4^ to 4.4 × 10^−4^). This corresponds to a rate of 5.3 × 10^−5^ per homologous gene copy that can act as a donor. We estimated that conversions occur in the intergenic spacer region between the 23S and 5S genes at a rate of 6.3 × 10^−5^ per genome (95% confidence interval: 1.6 × 10^−5^ to 1.6 × 10^−4^) or 1.1 × 10^−5^ per donor. The rate estimated for gene conversions in the spacer was similar to that for within the rRNA genes despite the greater sequence divergence in this region (Gurtler 1999, Liao 2000).

**Figure 3.**
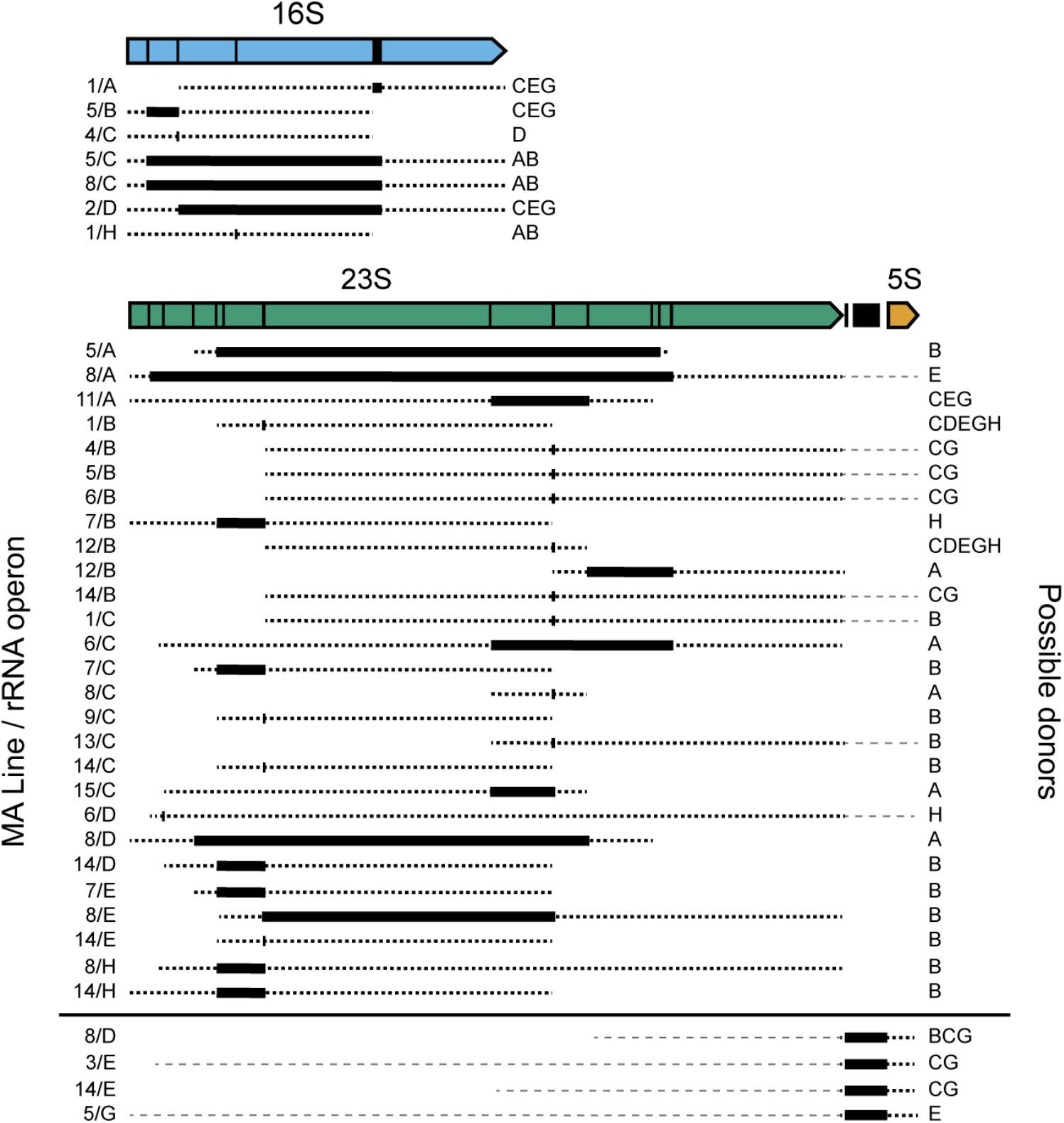
Gene conversions observed in rRNA operons during a mutation accumulation experiment. Bars in the genetic map indicate the locations of sequence differences between rRNA operons in the ancestral *E. coli* strain. In the remainder of the figure, boxes show the minimum extent of conversions and thick dashed lines show the maximum possible extent of conversions. Thick dashed lines show possible maximal extents that cross into the 23S-5S spacer. Possible donors are listed for the largest gene conversion events that could have resulted in only the observed sequence changes. Other donors may have been possible for smaller conversions. Events on each side of the 23S-5S spacer were considered separately so they were not included in the maximum sizes of these events when estimating rates (thin dashed lines).

A previous study of *E. coli* K-12 gene conversion rates that examined mutations that reverted the loss of a tRNA gene encoded in the spacer region between 16S and 23S genes found a mutant frequency of 6 × 10^−5^ per generation (Harvey and Hill 1990). This is roughly equivalent to the per-donor rate calculated here. Another study, using *E. coli* B rRNA operons, found a rate of 5 × 10^−9^ per donor per generation (Hashimoto et al. 2003). The significant discrepancy (10^4^ times less frequent) from our value of 5.3 × 10^−5^ is likely due to their use of counterselection against a *sacB*-*neo* cassette inserted into the *rrsB* gene as a means for recovering mutants. The insertion of this large (3825 bp) non-homologous region is expected to introduce a substantial barrier to conversion, as gene conversion rates fall sharply with reduced homology (Morris 2007).

The experiments presented here examine allelic gene conversion between copies of the rRNA operons. The *tufA* and *tufB* genes in *Salmonella typhimurium* have been used to quantify the ectopic conversion rate between homologous genes (Abdulkarim and Hughes, 2002). These genes share 98.9% sequence identity (1169/1182 bp) and experienced conversion at a frequency of 2 × 10^−8^ per generation, a rate approximately 1.6 × 10^4^ times less frequent than our per-donor allelic conversion rate. The rRNA operons in REL606 share between 99.2% and 100% sequence identity with each other, likely accounting for this difference.

We next examined whether certain rRNA operons were more likely to be converted than expected by chance (**Fig. 4a**). Of the 34 observed gene conversions within the 16S or 23S subunits, four were identified in *rrnA*, nine in *rrnB*, eleven in *rrnC*, four in *rrnD*, three in *rrnE*, and three in *rrnH*. No such gene conversions were identified in *rrnG*. Most operons showed no evidence of rate variation (adjusted *p* > 0.05), but *rrnC* was converted significantly more frequently than expected (adjusted *p* = 0.018). *rrnC* is the closest rRNA copy to the *E. coli* chromosome origin of replication (**Fig. 1a**), so it may exhibit an elevated conversion rate because it is the most likely to exist in multiple copies in one cell during DNA replication (Touchon et al. 2009). *rrnC* is also located in close proximity to *rrnA, rrnB, and rrnD*, which is expected to increase the chances that it will undergo homologous recombination (Hashimoto et al. 2003).

**Figure 4.**
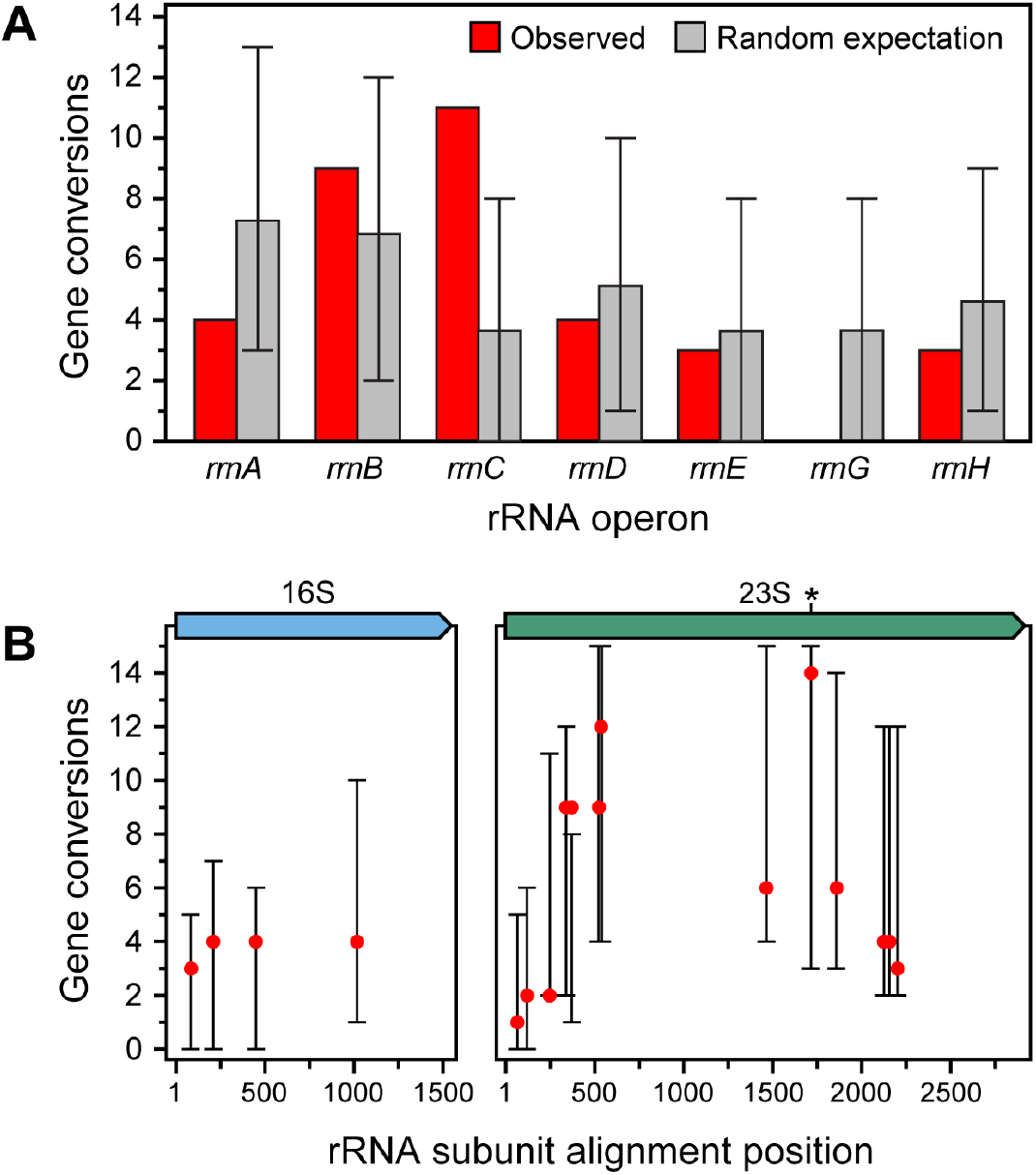
Distributions of conversion events changing the sequences of (A) rRNA operons and (B) heterologous sites within 16S and 23S rRNA sequences. The mean random expectation and error bars represent 95% confidence intervals estimated from 10,000 bootstrapping resamplings (see Methods). The location of the Chi-like site in the 23S subunit in *rrnA* is starred.

We also examined whether there was variation in the rates of conversions at certain sites within the 16S and 23S rRNA genes (**Fig. 4b**). We did not find strong evidence that any heterologous sites were converted significantly more or less frequently than expected (adjusted *p* > 0.05 for all sites). However, we did notice that sites at one position in the 23S subunit were converted fourteen times in the MA experiment and 2.3-fold more frequently than nearby sites approximately 200 bp away (**Fig. 4b**). The sequence at these sites is identical in six of the seven rRNA operons. In the seventh operon (operon A) this site contains a sequence (5′-GCTCGTGG-3′) that differs by one base from a canonical *E. coli* Chi site (5′-GCTGGTGG-3′). Similar Chi-like sites retain up to 40% of the activity of a consensus Chi site in promoting homologous recombination (Smith 2012), which suggests this sequence may be responsible for the somewhat elevated rate of gene conversions observed at this location.

Gene conversions maintain homogeneity between rRNA operons on long evolutionary timescales (Liao 2000, Hashimoto et al. 2003). Considered together, the 16S and 23S sequences in the seven rRNA operons have an average pairwise sequence identity of 99.578% in the ancestor of the MA experiment. The change in this identity ranged from +0.070% to −0.055% in the 15 MA lineages (**Fig. 5**), with no significant overall trend up or down (Wilcoxon signed rank test, *p* = 0.45). Sequence identity is only irreversibly ratcheted up by gene conversions when they completely eliminate an alternative allele from all copies of a gene, and the modest number of conversions that we observed was not enough to achieve this in nearly all cases. Therefore, this result underscores that homogenization via conversion is likely to only take place on longer timescales, particularly for sequences with many copies like the *E. coli* rRNA operons.

**Figure 5.**
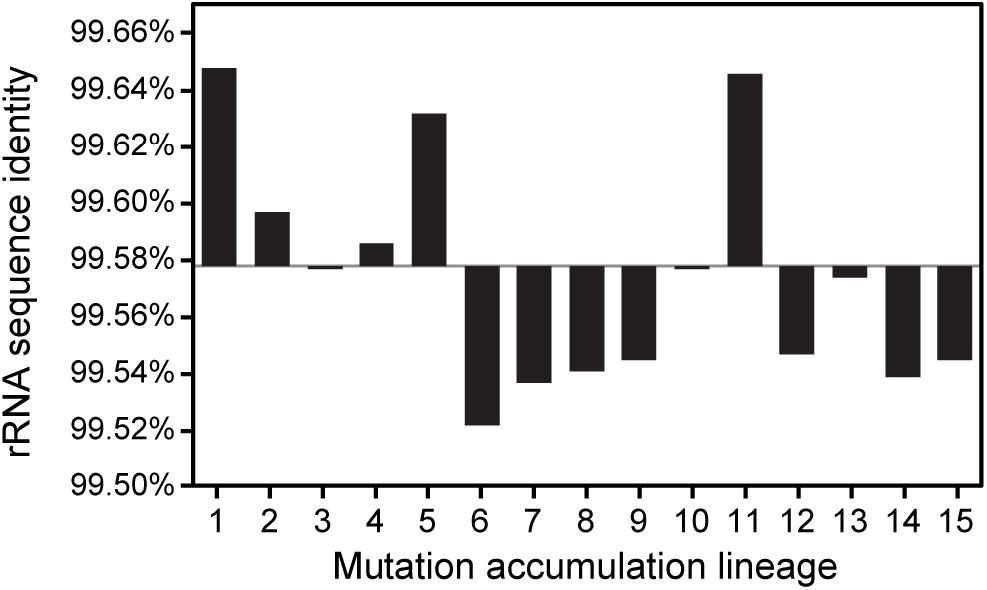
Evolution of average pairwise sequence identity in 23S and 16S rRNA subunits during the mutation accumulation experiment. The final value in each evolved lineage is depicted as a bar indicating the change from the 99.578% rRNA identity present in the ancestral *E. coli* strain.

The spontaneous rate of base substitution mutations in the *E. coli* is ~10^−10^ per base pair per generation (Wielgoss *et al.* 2011, Foster *et al.* 2012). It follows that the rate at which new base changes accumulate in the 23S and 16S genes in its seven rRNA operons is at most ~3 × 10^−6^ per genome per generation, and probably much less than this because many of these mutations will be deleterious. We found a gene conversion rate of ~3 × 10^−4^ per genome per generation, which is greater by two orders of magnitude. It is especially important to understand the relative balance and timescales of these mutational processes because rRNA sequences are used for phylogenetic reconstruction and metagenomic community profiling. Others have noted how heterogeneity in rRNA operons within a genome can complicate these analyses (Vetrovsky and Baldrian 2013, Ionescu *et al.* 2017) and how gene conversions can obscure evidence of rRNA horizontal gene transfer (Tian *et al.* 2015). Conversion rates also influence the chances that paralogs can evolve sequence diversity and stably maintain different functions (Teshima and Innan 2008). Additionally, gene conversions have been observed in cultures through routine laboratory use, including one conversion in the *rrlH* gene of REL606 (Studier *et al.* 2009). Our measurements of *E. coli* rRNA gene conversion rates enable more accurate modeling of these processes and improve our overall understanding of bacterial genome evolution.

## Acknowledgments

This work was supported by a Welch Foundation grant (F-1979) and a US Army Research Office grant (W911NF-12-1-0390).

